# First chromosome-level genome assembly of the colonial tunicate *Botryllus schlosseri*

**DOI:** 10.1101/2024.05.29.594498

**Authors:** Olivier De Thier, Mohammed M.Tawfeeq, Roland Faure, Marie Lebel, Philippe Dru, Simon Blanchoud, Alexandre Alié, Federico D. Brown, Jean-François Flot, Stefano Tiozzo

## Abstract

*Botryllus schlosseri* (Tunicata) is a colonial chordate that has long been studied for its multiple developmental pathways and regenerative abilities and its genetically determined allorecognition system based on a polymorphic locus that controls chimerism and cell parasitism. We present the first chromosome-level genome assembly from an isogenic colony of *B. schlosseri* clade A1 using a mix of long and short reads scaffolded using Hi-C. This haploid assembly spans 533 Mb, of which 96% are found in 16 chromosome-scale scaffolds. With a BUSCO completeness of 91.2%, this complete and contiguous *B. schlosseri* genome assembly provides a valuable genomic resource for the scientific community and lays the foundation for future investigations into the molecular mechanisms underlying coloniality, regeneration, histocompatibility, and the immune system in tunicates.

## Introduction

In the sub-phylum Tunicata, which is the sister group of vertebrates (1), colonial species can reproduce both sexually and asexually via different forms of budding. Asexually generated individuals remain generally connected, hence the name “colonial”. Through budding, a new functional body arises from adult somatic cells and tissues. Budding ontogeny varies among species, involving non-homologous tissues and cells (2). Despite differences in budding modes and whether development occurs asexually or sexually via embryogenesis, the Bauplan of an adult tunicate remains largely conserved across the entire sub-phylum (3).

Like many other colonial tunicates, *Botryllus schlosseri* can generate a functional adult body via three distinct developmental pathways. The first pathway involves sexual reproduction, in which a fertilized egg passes through a larval stage and develops into the first individual of the future colony. The second pathway involves asexual propagation, in which the founder zooid reproduces continuously through a process called palleal (or peribranchial) budding. The outcome is a colony of up to hundreds of zooids connected by an extracorporeal network of blood vessels and embedded in a cellulose-based extracellular matrix called “tunic” (4). Finally, if all zooids and buds are removed from a colony of *B. schlosseri*, new buds can develop from the extracorporeal vascular system through a regenerative process known as vascular budding (5). The regenerating zooids propagate asexually and eventually reform a new colony (6, 7).

Zooids belonging to the same colony are essentially all clones. However, in the wild, nearby colonies frequently come in contact and fuse, forming chimeras. Through *Botryllus* chimerism, multiple pools of circulating cells from different genotypes come into contact throughout their lifetime. All of these pools contribute to sexual, asexual, and regenerative development (8–10). During chimerism, cells from the donor can also completely replace the cells of the host in both the germline and/or the somatic tissues. These processes are known as somatic cell parasitism or germ cell parasitism depending on whether somatic cells or germ line cells are concerned (10–12). Therefore, the mechanisms of cell parasitism and the dynamic nature of budding imply that zooids in chimeric colonies may not always be clonemates.

*Botryllus schlosseri* was introduced in laboratories more than half a century ago (13), mainly as a model to study asexual development, regeneration (14), allorecognition and chimerism (15, 16). Over the last decades, a scientific community has taken shape and several research groups have made significant efforts to improve breeding conditions (17– 19) and to develop and adapt imaging and molecular biology techniques (7, 8, 20). Several anatomical descriptions and staging methods have been proposed (4, 21) and various stage-specific and tissue-specific transcriptomic databases have been produced (7, 22–26). In 2013, a draft genome of *B. schlosseri* become available (27). Yet, this published assembly lacked completeness and contiguity and did not meet the current assembly standards (28). In the present study, we provide high-quality, chromosome-level, haploid assembly of the *Botryllus schlosseri* genome. This new assembly represents a handy and usable resource for exploring the nature of developmental and regenerative mechanisms that characterize this species, to elucide its complex allorecognition, chimerism, and cell parasitism, and in general to better understand the biology of colonial chordates.

## Results and Discussion

### Genome sequencing and assembling

To generate a new, high-quality genome assembly, we sequenced gDNA extracted from a laboratory-reared non-chimeric colony derived from a single zygote, referred to as clone E*, from which we obtained a combination of 489 million Illumina (short) paired-end 150 bp reads, 2.4 million PacBio HiFi (long) reads with a N50 length of ∼9.5 kb (max length of ∼50 kb) and 7.4 million ONT (long) reads with a N50 length of ∼10.5 kb (max length of ∼205 kb) (Table 1).

**Table 1.** Sequencing technology applied to sequence *B. schlosseri* genome (clone E*) and relative read statistics.

| Technology | Tot. size (Gbp) | Number of reads | N50 (bp) | Coverage |
| --- | --- | --- | --- | --- |
| Illumina | 73.2 | 488,906,094 | 150 | 146 |
| Illumina Hi-C | 15.9 | 106,488,252 | 150 | 32 |
| PacBio HiFi (round 1) | 7.9 | 1,170,137 | 8,711 | 16 |
| PacBio HiFi (round 2) | 10.8 | 1,218,052 | 10,151 | 22 |
| ONT (R9.4.1) | 58.9 | 10,888,103 | 10,320 | 118 |

Based on k-mer analyses, the genome size was estimated to be around 500 Mbp (∼444 Mbp using a maximum k-mer count of 10,000 and ∼515 Mbp using a maximum k-mer count of 10,000,000) with a heterozygosity of 3.63% (Figure S1), whereas the Feulgen estimate was *∼*499 Mbp (using 1 pg = 978 Mbp) (Figure S2), which is less than a previous cytofluorimetric-based estimation of 725 Mb (29) and than the first genome assembly obtained by Voskoboynik *et al*. (27), which had a size of 580 Mbp.

An initial purged primary assembly was obtained using hifiasm (30); it had a size of 570 Mbp and comprised 930 contigs with an N50 length of 4.9 Mbp. In that assembly, BlobToolKit analyses found 435 contigs with a total size of 36 Mbp corresponding to putative contaminations that were subsequently removed. Among these 36 Mbp, around half belonged to members of the bacteria phylum Pseudomonadota (Figure S4). The entire mitochondrial DNA was retrieved in a single contig of length 32,493. BLASTN comparison between this contig and the previously published mitochondrial genome of *B. schlosseri* (27) shown that this contig contained duplications and misassemblies, hence we manually removed from the assembly. The remaining contigs were corrected using CRAQ (31), which detects and break misassembled contigs; this raised the total number of contigs in the assembly from 477 to 516. We then performed Hi-C scaffolding using YaHS (32), which reduced the number of contigs to 256, before running CRAQ again on the scaffolded assembly (this time, 4 misassembled contigs were detected and broken). Finally, a manual curation was performed, resulting in an assembly made up of 16 major scaffolds containing around 96% (513 Mbp) of the total sequence length (533 Mbp) (Table 2, Figures 1 and 2). The full assembly pipeline is summarized in Figure 4 and detailed in the Material and Methods section.

**Table 2.** Assembly statistics for all the scaffolds and for the 16 longest ones.

| Measure | All scaffolds | 16 longest scaffolds |
| --- | --- | --- |
| Length (Mbp) | 533 | 513 |
| No. of scaffolds | 254 | 16 |
| N50 (Mbp) | 30 | 31 |
| GC (%) | 40.52 | 40.46 |
| No. of annotated genes | 22,275 | 21,676 |
| BUSCO Complete (Single, Duplicated) | 91.2% (90.0%, 1.2%) | 91% (90.1%, 0.9%) |
| BUSCO Fragmented | 3.4% | 3.2% |
| BUSCO Missing | 5.4% | 5.8% |

**Fig. 1.**
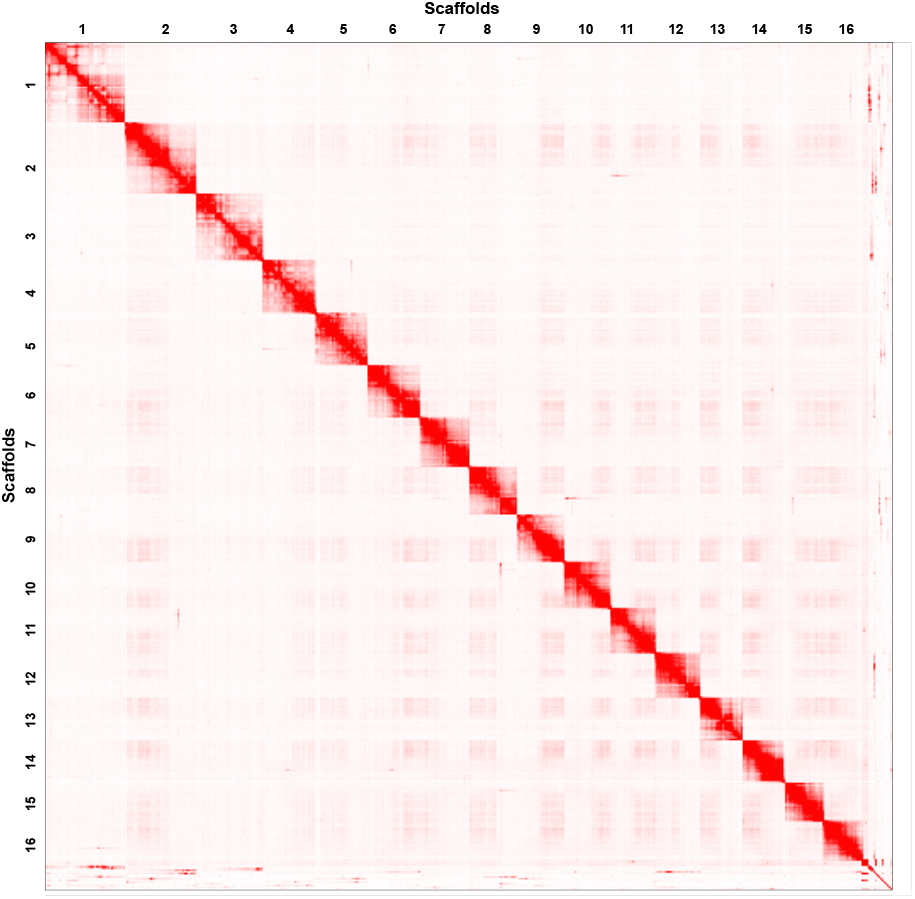
Hi-C heatmap of the assembly of the genome of *Botryllus schlosseri* showing sixteen chromosome-scale scaffolds.

**Fig. 2.**
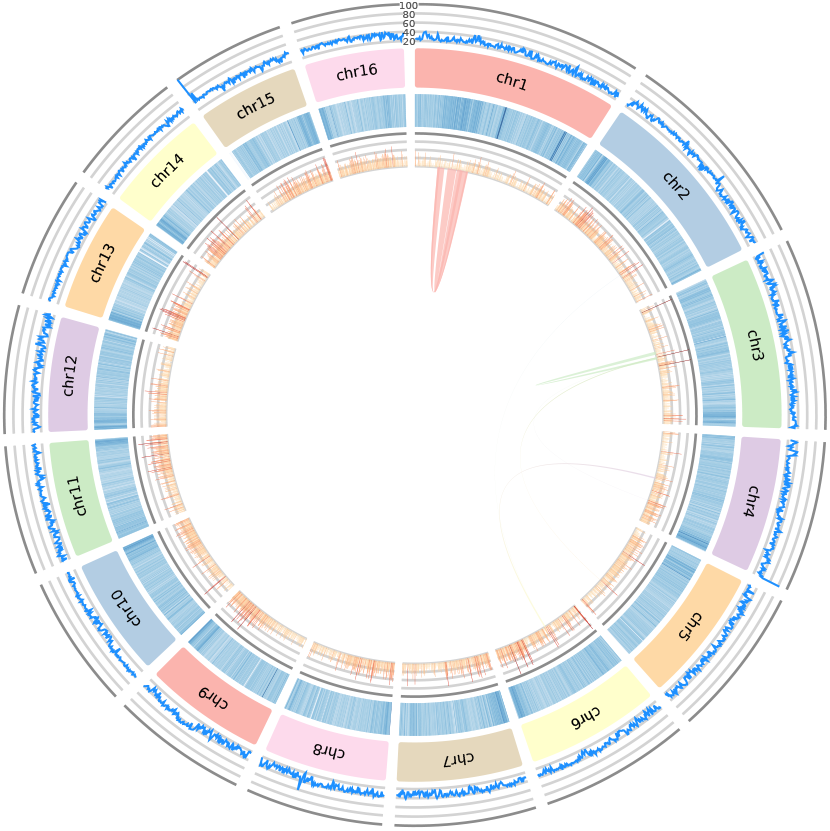
Circos plot of the distribution of several genomic characteristics along the 16 longest scaffolds represented as chromosomes made with AccuSyn (33). From the inner to the outer circles, are represented: the syntenic blocks retrieved with MCScanX (34); the histogram of the gene density; a heatmap of the presence of repetitive elements; the reference chromosomes in clockwise orientation and the read coverage using the HiFi reads.

The completeness of our assembly was then assessed using the Benchmarking Universal Single-Copy Orthologs (BUSCO) tool (35), which returned a genome completeness of 91.2% (including 1.2% of duplicated marker genes), compared to 74.4% (including 23.7% of duplicated genes) for the assembly of Voskoboynik *et al*. (Figure 3). This high duplication score of the previously available assembly indicates that their larger assembly size (580 Mbp vs. 533 Mbp) was caused by incompletely collapsed haplotypes (36). Synteny analysis performed with MCScanX (34) highlighted the presence of two large-scale genomic palindromes located within Bs1 (Figure 2). To find out whether these palindromes may result from assembly artifacts (37), we checked the localisation of the duplicated BUSCO genes along the chromosomes and did another run of CRAQ using this time ONT as long reads (with double coverage compared with the HiFi reads used in the previous rounds). There was no significant difference in the number of duplicated BUSCO genes within Bs1 compared to the others, and CRAQ did not detect structural errors in this scaffold either. These observations suggest that the palindromes observed in Bs1 are real, with potential biological significance that will require further investigation.

**Fig. 3.**
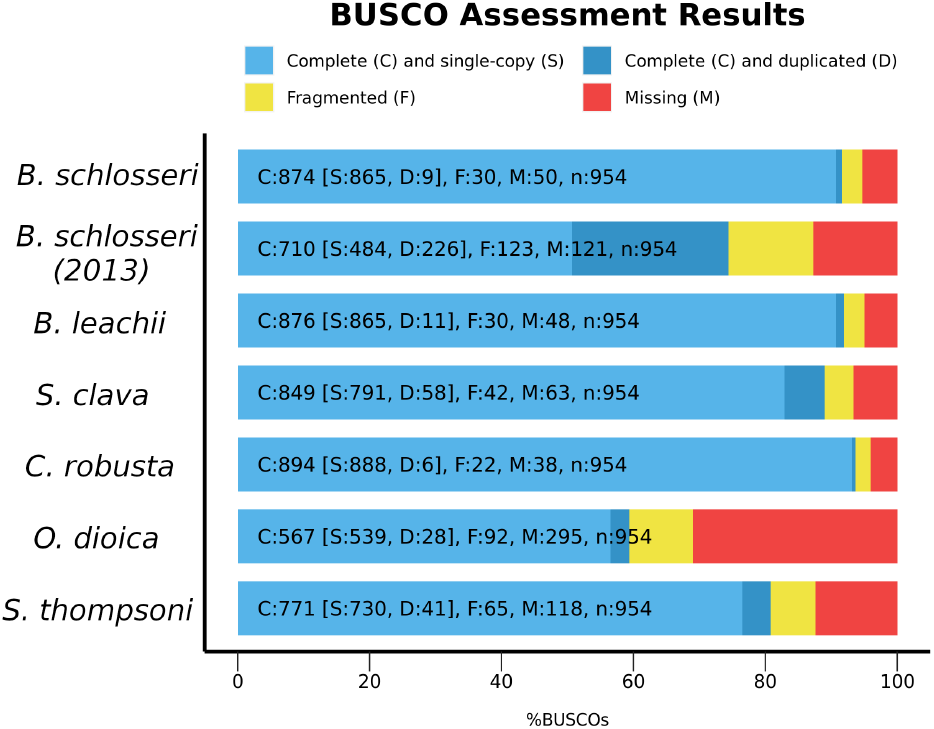
Orthology assignemnt in previous tunicate genome projects. Proportion of BUSCO genes detected or missed in the new genome assembly of *B. schlosseri* compared to the existing assembly (*B. schlosseri (2013)* (27)) and reference ones.

**Fig. 4.**
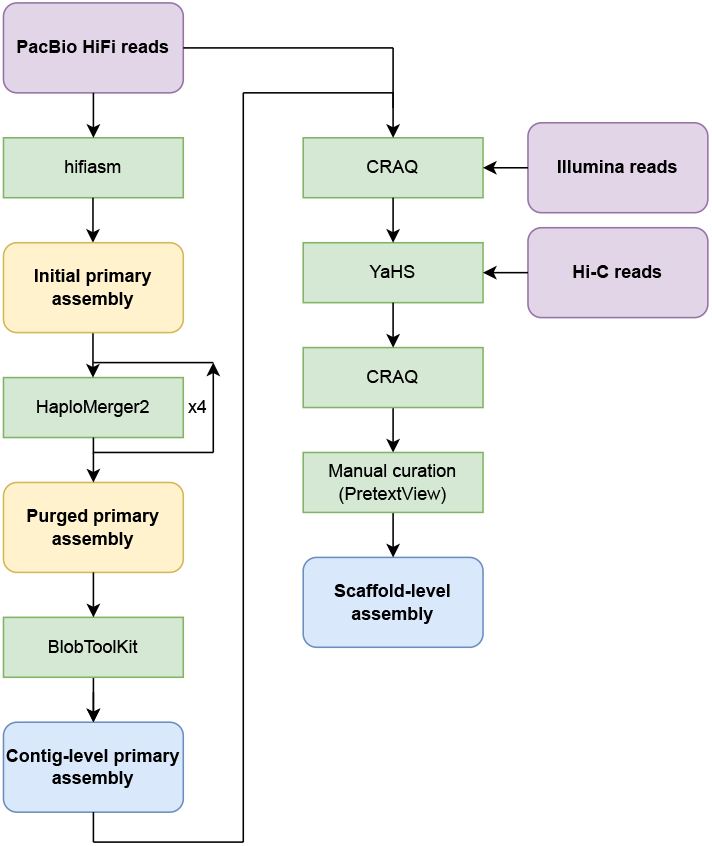
Assembly pipeline (see Material and Methods.)

### Structural and functional annotation

Before gene annotation, RepeatMasker, using a *de novo* repeat library created by RepeatModeler, estimated that around 63% of the *B. schlosseri* genome assembly consisted of repetitive elements, which is close to the 65% of repeats fond in the previously published assembly (27). Most of these were interspersed repeats (see Table 3). *Ab initio* genome annotation using the BRAKER3 pipeline (38) initially predicted 16,966 coding genes, after which refinement using the PASA pipeline (39, 40) finally retrieved 22,275 genes coding for 30,813 proteins (see Table 4). This number is significantly lower than originally predicted for *B. schlosseri* (38,730 predicted genes (27)), probably due to the incomplete collapse of the previous assembly. In terms of completeness of the annotation, BUSCO retrieved 92.4% complete (79.8% single, 12.6% duplicated) and 1.8% fragmented metazoan genes, whereas 92.2% complete (91.2% single, 1% duplicated) and 1.8% fragmented of those genes were retrieved when filtered to only keep their longest isoform. This is consistent with the results obtained by running BUSCO directly on the scaffold sequences.

**Table 3.** Classes of repeats in the *Botryllus schlosseri* genome. RepeatMasker summary table output for the haploid genome of *Botryllus schlosseri* showing the different classes in percentages of identified repeats.

| Repeat class | Percent of genome |
| --- | --- |
| **LINEs** | **4.52%** |
| LINE1 | 0.15% |
| LINE2 | 2.06% |
| **LTR elements** | **1.34%** |
| **DNA elements** | **7.24%** |
| hAT-Charlie | 2.96% |
| TcMar-Tigger | 0.01% |
| **Unclassified** | **46.03%** |
| **Total interspersed repeats** | **59.12%** |
| **Simple repeats** | **3.94%** |
| **Low complexity** | **0.02%** |
| **Total** | **63.09%** |

**Table 4.** Gene predictions and annotation statistics, including initial gene models predicted from Braker3 pipeline.

| Type | Number | Mean size (bp) | % genome |
| --- | --- | --- | --- |
| Gene | 22275 | 8566.13 | 35.78 |
| mRNA | 30813 | 10576.62 | 61.11 |
| cds | 237200 | 199.16 | 8.86 |
| Exon | 241815 | 289.83 | 13.14 |
| 5' UTR | 21386 | 432.29 | 1.73 |
| 3' UTR | 20985 | 648.00 | 2.55 |
| Total | 574474 | 1143.44 | 123.18 |

The functional annotation and orthology assignment (41), coupled with annotation of protein domains, motifs, and functional sites (42, 43), were into gff3 and Genbank files (available upon publication). KEGG route-mapping assigned 7,221 genes over the annotated entries and distributed them across 21 KEGG categories (Figure 5). Among them, the most prevalent ones include different KEGG hierarchies dealing with genetic information processing (2449/7219, 22,92%), such as DNA replication, repair, recombination, transcription, translation and regulation of gene expression; signaling and cellular processes (886/7219, 12,27%); and environmental information processing (674/7219, 8,64%) such as various cellular processes and signaling pathways involved in sensing, transducing (i.e. MAPK signaling, PI3K-Akt signaling and cAMP signaling), responding to external signals (i.e. G-protein coupled receptors, receptor tyrosine kinases, and cytokine receptors), intracellular communication and cell motility. The KEGG annotations provided for *B. schlosseri* are consistent and coherent with the functional annotation of the published complete genomes of other ascidian tunicates such as *Styela plicata, Ciona robusta* and *Oikopleura dioica* (Figure S5).

**Fig. 5.**
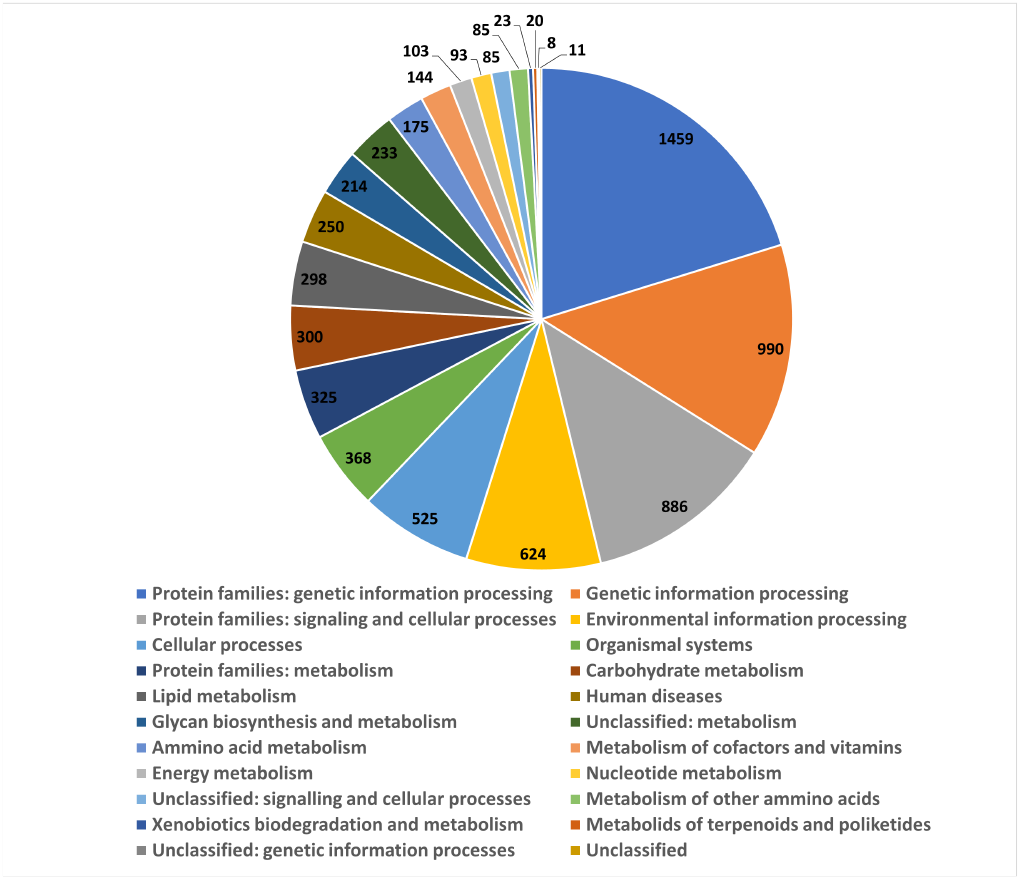
Pie chart of the assignation of the annotated genes of *Botryllus schlosseri* to KEGG functional categories using BlastKOALA (44)

### Synteny analyses

To assess macrosynteny conservation between *B. schlosseri* and other tunicates, we compared our assembly with the three currently available tunicate genomes assembled at chromosome-level, namely *Styela clava* (45) from the same order as *Botryllus* (Stolidobranchia), *Ciona robusta* (46), from a different order (Phlebobranchia), and *Oikopleura dioica* (47) from a different class of tunicates (Appendicularia) (48). We used 17 groups of orthologous genes identified by Simakov *et al*. as ancestral chordate linkage groups (CLGs) (49). These represent groups of genes that have remained physically linked since the divergence of the olfactores (= vertebrates + tunicates) and cephalochordate lineages. The analyses generated using Oxford dot plots (50) revealed units of conserved synteny (Figure 6), represented in Figure S6 by dense rectangular blocks of dots. This physical linkage was generally conserved among *B. schlosseri, S. clava* and *C. robusta*, yet the random distribution of ortholog pairs within blocks suggests that the gene order within syntenic units has become scrambled, resulting in a loss of colinearity. The comparison with *Oikopleura dioica* showed a complete loss of both synteny and colinearity due to chromosome fission and extreme mixing. The latter result is in line with the very long and fast-evolving branch of the Appendicularia when compared to the other tunicates (48) as well as an extreme genome scrambling rate of Appendicularia compared to other tunicates and mammals (51). The same analyses using a set of 29 linkage groups generally conserved among bilaterians (52) yielded similar results (not shown).

**Fig. 6.**
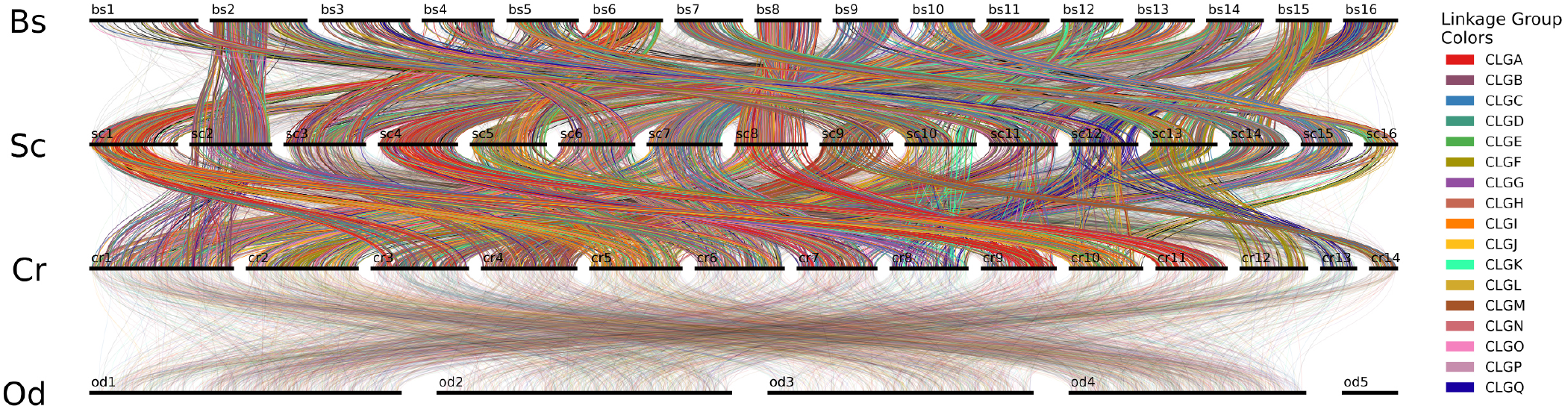
Synteny conservation of chordate linkage groups (CLGs) (49, 52) among *Botryllus schlosseri* (Bs), *Styela clava* (Sc), *Ciona robusta* (Cr) and *Oikopleura dioica* (Od).

### Hox gene analyses

Hox genes are a subset of homeobox genes that play important developmental roles in the specification of body segments along the anterior-posterior axis. Their arrangement into a syntenic cluster colinear with gene expression is conserved across Bilateria, with some exceptions (53). In the current assembly, we retrieved ten *B. schlosseri* Hox genes, which is in line with draft genomes of other ascidian tunicates (54). Orthology of *B. schlosseri* Hox genes was assessed using phylogenetic analyses as in Sekigami *et al*. (55), based on Hox tree topology among the tunicates *Ciona robusta* and *Halocytnthia roretzi*, the cephalochordate *Branchiostoma lanceaolatum* and three vertebrate species. The names of the *B. schlosseri* Hox genes were assigned based on their proximity to the ones of *C. robusta* (Figure S7). However, most branches were not statistically robust, and therefore including more tunicates as well as vertebrate species will be necessary to resolve the complex evolution of the Hox gene cluster across tunicates (54).

Although Hox genes are colinear between cephalochordates and vertebrates, it is not the case for tunicates (56). In the tunicate species studied thus far, the Hox clusters exhibit divergences in terms of colinearity and synteny relative to the ancestral chordate cluster (54). In contrast to previous data (27, 57), our new assembly revealed that *B. schlosseri*’s Hox genes are not scattered. Instead, eight of them were clustered on the second largest scaffold (Bs2), whereas two other ones are found on the 15th largest scaffold (Bs15). Comparison with two tunicate ascidians, belonging to the same (*H. roretzi* (55)) and a different (*C. robusta* (46)) order, revealed partially conserved synteny as well as inversions and transpositions across the three species (Figure 7). These observations agree with the general trend of synteny conservation despite loss of colinearity observed for CLGs (49) and are also consistent with the phylogenetic relationships among the species sequenced (2, 48).

**Fig. 7.**
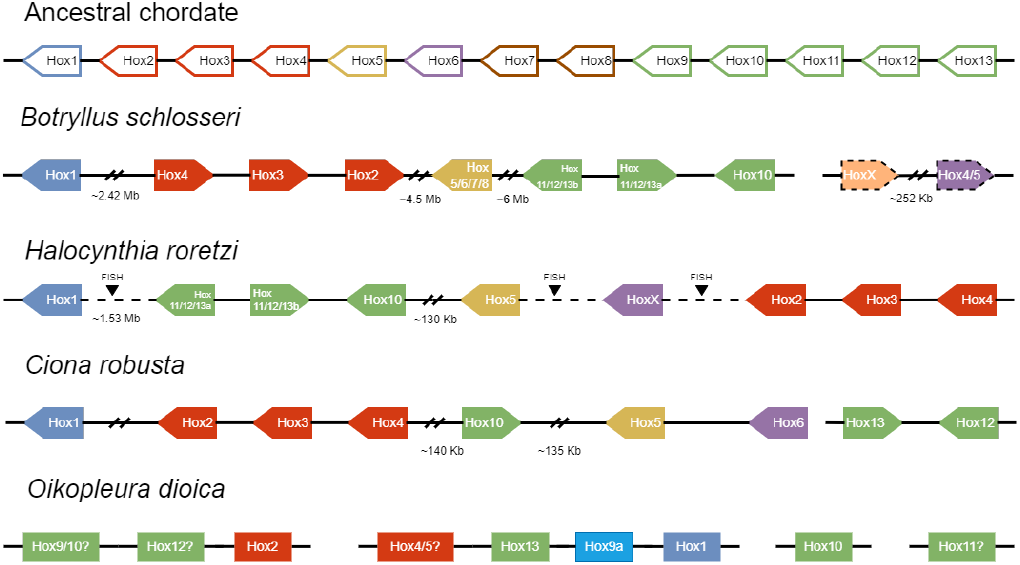
Representation of the Hox genes retrieved in the new assembly of *B. schlosseri* compared to the supposed original single Hox cluster of the chordate ancestor and other tunicates. Linked genes (present on the same scaffold) are connected by a solid line while a dashed line is used when the linkage has been deduced with another method. When known, the transcription orientation is given by an arrow-shaped rectangle, which is surrounded by a dashed line when the Hox gene was retrieved with low confidence.

## Conclusion

Our novel, high-quality chromosome-level assembly of the *Botryllus schlosseri* genome is suitable for analyses that require a comprehensive and highly contiguous genomic reference. This new resource will enable better comparative genomic studies and will serve as a basis for broader investigations into the molecular mechanisms that control developmental processes and immune responses in tunicates.

## Materials and Methods

### Sampling, DNA isolation, and sequencing

Isogenic colonies of *Botryllus schlosseri* were raised on glass slides in the marine-culture system described in Langenbacher *et al*. (20). Genomic DNA was extracted from the colony labeled E* using Qiagen’s MagAttract HMW DNA Kit (67563). Libraries were prepared and sequenced performed at Novogene (Cambridge, UK) for Illumina 2x150 bp paired-end (PE) reads, at the Leiden Genome Technology Center (Leiden, Netherlands) for HiFi PacBio long reads and at UCAGenomix (Valbonne, France) for Oxford Nanopore (ONT) long reads (on a FLO-PRO002 flow cell with R9.4.1 pore proteins, using the SQK-LSK109 ligation sequencing kit). Nanopore base calling was performed using Guppy v3.2.10. A Hi-C library was prepared using the Arima High Coverage HiC Kit (A410110) followed by the Arima HiC+ Kit (A510008, A303011) and sequenced using Illumina (2x150 bp).

### RNA-seq data

Illumina PE RNA-seq reads were retrieved from the published datasets of Rodriguez *et al*. (22) and Ricci *et al*. (7, 24), for a total of about 239 Gbp.

### Data preprocessing

PacBio HiFi reads were processed with HiFiAdapterFilt v2.0.1 (58) to remove adapter sequences, while Porechop v0.2.4 (https://github.com/rrwick/Porechop) was used to trim basic adapters from ONT reads. For Illumina reads, quality trimming and adapter clipping were performed using Trimmomatic v0.39 (59) while quality check, prior to and after trimming, was done using FastQC v0.11.5 (60).

### Genome size estimation

The genome size of colony E* was measured using the improved Feulgen protocol of M.Tawfeeq et al. (preprint in prep.). In brief, the protocol steps included: chopping the tissue into tiny pieces using a sterilized razor blade with a few drops of 40% glacial acetic then leaving it for 48 hours in the dark; immersing the processed slides into th fixation reagent (85:10:5 volumes of methanol:formaline:acetic acid); then hydrolysing them (using hydrochloric acid) and staining them (using Schiff’s reagent). We used three standards of known C-values: *Periplaneta americana* (3.41 pg) (61), *Lasius niger* (0.33 pg) (JF Flot, unpublished data), and *Ambystoma mexicanum* (32 pg) (62). A digital camera (5 megapixels) mounted on a compound microscope was used for imaging the slides, and nuclei measurements were performed using ImageJ (63).

A genome size estimation based on the k-mer spectrum of the Illumina reads was also performed using KMC v3.2.1 (64) and the GenomeScope2.0 (65) web server, with a k-mer size of 21 and maximum k-mer counts of 10,000 and 10,000,000.

### Genome assembly

First, the PacBio HiFi reads were assembled into contigs using hifiasm v0.19.5-r592 (30) with the haplotype purging option disabled (option -l0 with hifiasm in HiFi-only Assembly mode). Second, uncollapsed haplotypes were purged using multiple rounds of HaploMerger2 (release 20180603) (66) until the BUSCO duplication score stabilized. Third, non-metazoan contigs were identified and removed from the assemblies using BlobToolKit v4.1.5 (67). To his aim, contigs were aligned to the NCBI nucleotide database (accessed 2023 March 18) using BLAST+ (68) with the blastn command, and a also to the UniProt reference proteome database (accessed in 2023 March 23) using DIAMOND v2.1.6 (69); contig HiFi coverage depth was computed using minimap2 v2.24-r1122 (70). Using the “bestsumorder” rule of BlobToolKit, only the contigs assigned to the taxon “Chordata” or without a match (“no-hit”) were kept. Finally, a BLASTN search for fragments of the mitochondrial genome among the contigs was performed using the published complete mitochondrial genome of *B. schlosseri* (RefSeq NC_021463.1) (27).

To scaffold the assemblies, PacBio HiFi and Illumina reads were first mapped to the assemblies with minimap2 and putative misjoined regions were identified and automatically split using CRAQ v1.0.9 (31) with default parameters except for the addition of –break. Hi-C reads were subsequently mapped to the output of CRAQ using the Arima Genomics mapping pipeline script arima_mapping_pipeline.sh (71) (https://github.com/ArimaGenomics/mapping_pipeline), and YaHS v1.2 (32) was run with default parameters to scaffold the assemblies. CRAQ was then applied to the results, and finally the scaffolds were manually curated using PretextMap v0.1.9 (72) and PretextView v0.2.5 (73). Metrics for the assemblies were computed with SeqKit v2.3.0 (74) (parameter stats -a). The quality and completeness were checked using KAT v2.4.2 (75) on k-mers from both PacBio HiFi and Illumina reads, and BUSCO v5.4.4 (76) (using the -m genome mode) with the metazoa_odb10 dataset.

### Genome annotation

Repetitive elements were identified using RepeatModeler and RepeatMasker pipeline. A *de novo* repeat library was generated using RepeatModeler2 v2.0.3 (77) and used as input for RepeatMasker v4.0.6 (78) to detect, classify and soft-mask repeats in the genomic sequences. RNA-seq reads were aligned to the soft-masked assemblies using STAR v2.7.10b (default options) (79). Based on the aligned transcripts, a list of proteins from OrthoDB v11 (80) for Metazoa (https://bioinf.uni-greifswald.de/bioinf/partitioned_odb11/) as extrinsic evidence and the soft-masked assemblies, gene prediction and annotation was done using the BRAKER3 v3.06 pipeline for RNA-Seq and protein data without training or gene prediction with untranslated regions (UTRs) parameters (38, 81–93). A refinement of the initial BRAKER3 structural annotation and the addition of UTRs were then performed with an implementation the PASA pipeline v2.4.1 (39) in conjugation with EVidenceModeler (EVM) v2.1.0 (40). A third of the RNA-seq reads of the Rodriguez *et al*. (2014) transcriptome (22) were aligned again to the assemblies and their BRAKER3 annotation using STAR v2.7.10b (MAX_INTRON_SIZE=20000), then (79) and assembled with StringTie v2.2.1 (94) using the BRAKER3 annotation as a reference. PASA_alignment_assembly was then run with the transcripts assembled by StringTie and independently with Trinity assemblies of the Rodriguez *et al*. (2014) RNAseq (22), Ricci *et al*. (2016) RNAseq (24) and Ricci *et al*. (2022) RNAseq (7). TransDecoder (95) was run within PASA to identify coding sequences within the assembled transcripts. A consensus annotation of coding sequences (CDSs) was found by EVM by leveraging both the transcripts and coding sequences identified for each RNAseq by PASA (evidence weights: 1 for BRAKER3 input, 5 for PASA transcripts and TransDecoder CDSs). The UTRs and isoforms were added by running PASA_genome_annotation sequentially with each previously generated PASA database (using EVM output as the first reference, then the output of the previous PASA_genome_annotation run). Functional annotation was performed starting from the structural annotation obtained with BRAKER3-PASA pipeline. Eggnog-mapper (41, 96) pipeline combined with Interproscan (42, 43) was used for orthology-based annotation (nr, KEGG, GO terms) and for protein domains prediction respectively. Both approaches were used as input for the Funannotate v1.8.15 pipeline (97), yielding a gff3 and a Genbank file with functional annotations (available upon request).

### Comparative genomics analyses

The genome assemblies and annotations for the comparison with other tunicate species were retrieved from ANISEED (98) for *Botrylloides leachii, Ciona robusta*, and for the first assembly of *Botryllus schlosseri*, while *Oikopleura dioica* originates from (47), *Salpa thompsoni* from (99) and *Styela clava* from (45). Macrosynteny analyses were performed using the odp tool (50). For each species, analyses were based on the longest protein isoforms generated from their annotation file using AGAT’s (100) scripts agat_sp_keep_longest_isoform.pl and agat_sp_extract_sequences.pl -p.

### Phylogenetic analyses of Hox genes

Phylogenetic analyses of *Botryllus schlosseri* Hox genes were performed using sequences retrieved from Sekigami *et al*. (55). First, the sequences were aligned using MUSCLE (101) as implemented in AliView (102), then IQ-TREE 2 (103) was used to build a Maximum Likelihood phylogeny with the best-fit model JTT+R6 (104, 105) selected by ModelFinder (106) following the Bayesian information criterion (107) and with 10,000 ultrafast bootstrap replicates (108).

## ACKNOWLEDGEMENTS

We would like to thank EMBRC-France and in particular Laurent Gilletta for isolating isogenic colonies and maintaining the aquaculture system. This work was supported by ANR (ANR-14-CE02-0019-01), INSB-DBM, Sorbonne Universite AAP Emergence 2021 and FAPESP 15/50164-5 & 19/06927-5.

## Supplementary Material

**Fig. S1.**
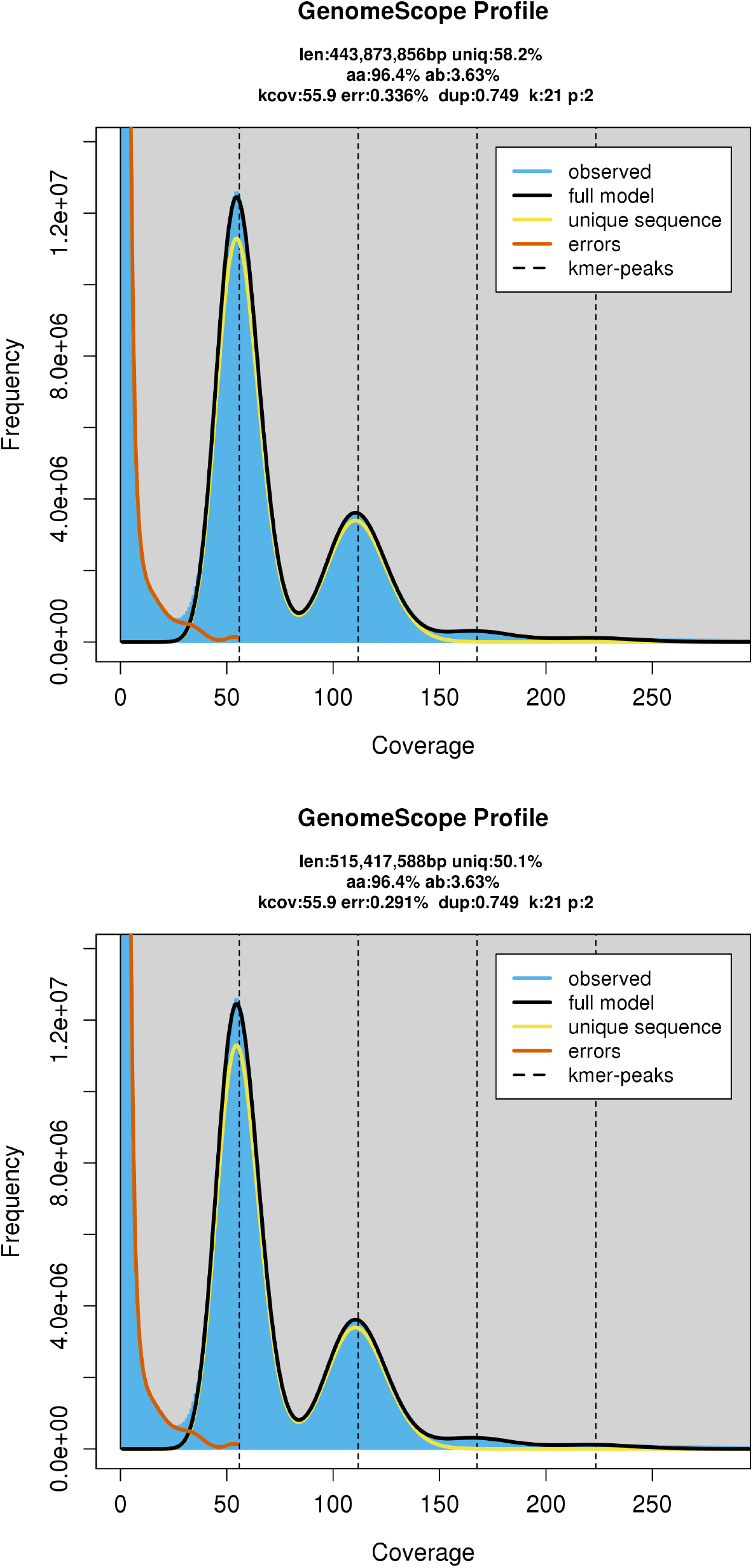
Genomescope2.0 results obtained with the Illumina reads and using 21-mers with a maximum counts of 10,000 (top) and 10,000,000 (bottom).

**Fig. S2.**
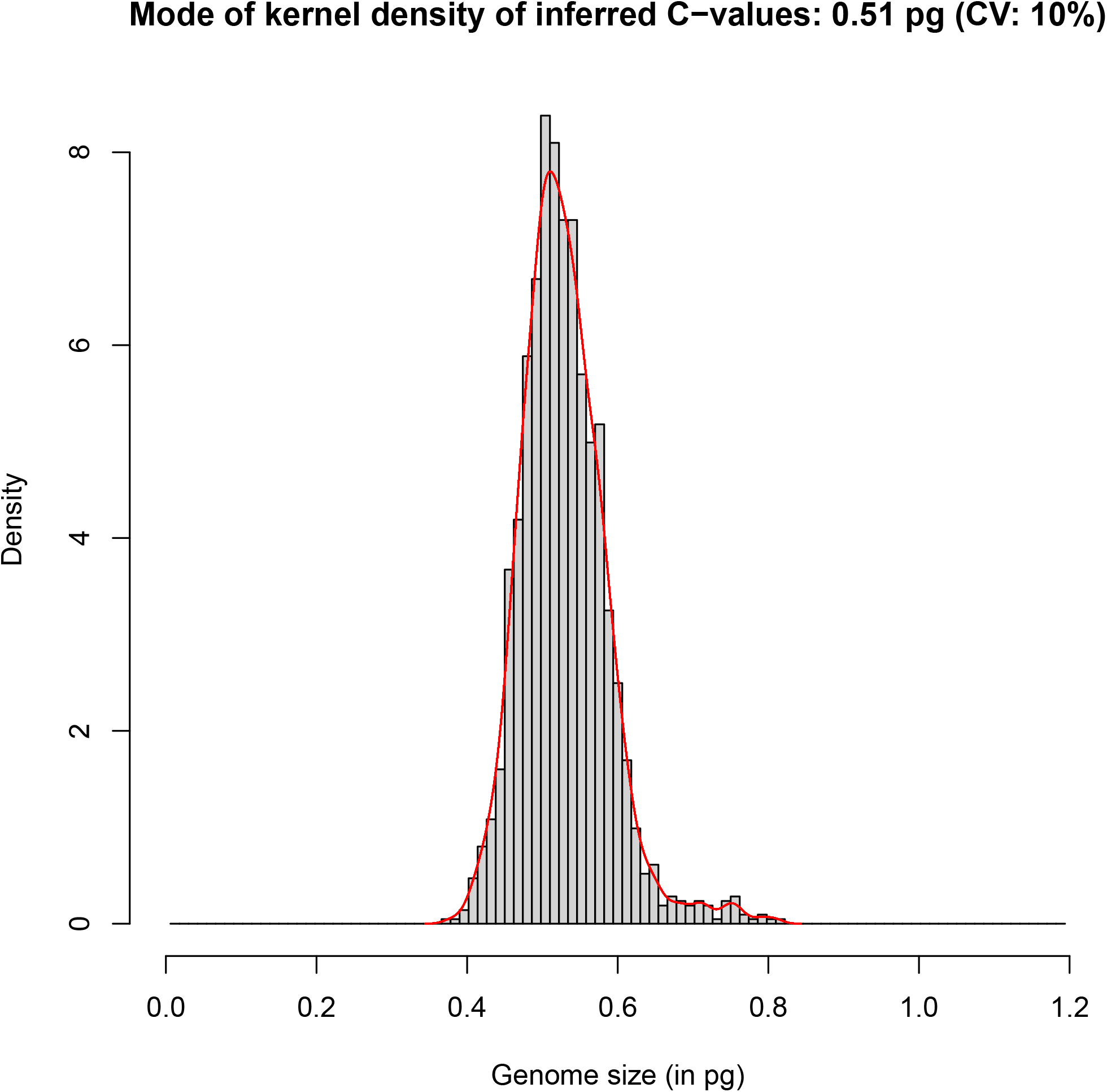
Genome size of *Botryllus schlosseri* measured using Feulgen microphotodensitometry.

**Fig. S3.**
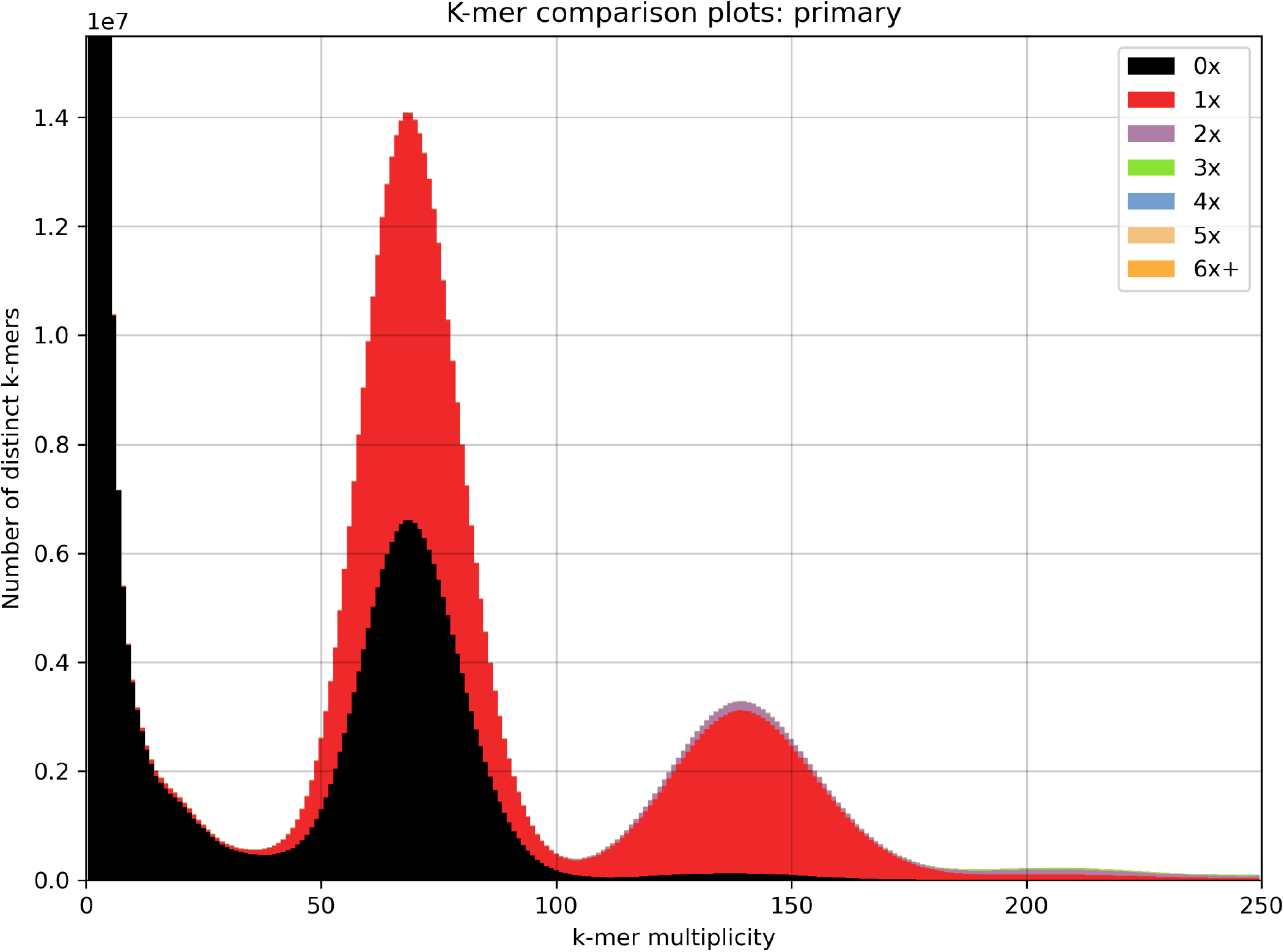
Output of the KAT comp tool comparing k-mers generated from a concatenation of the Illumina and HiFi reads to those generated from the primary assembly of *B. schlosseri*.

**Fig. S4.**
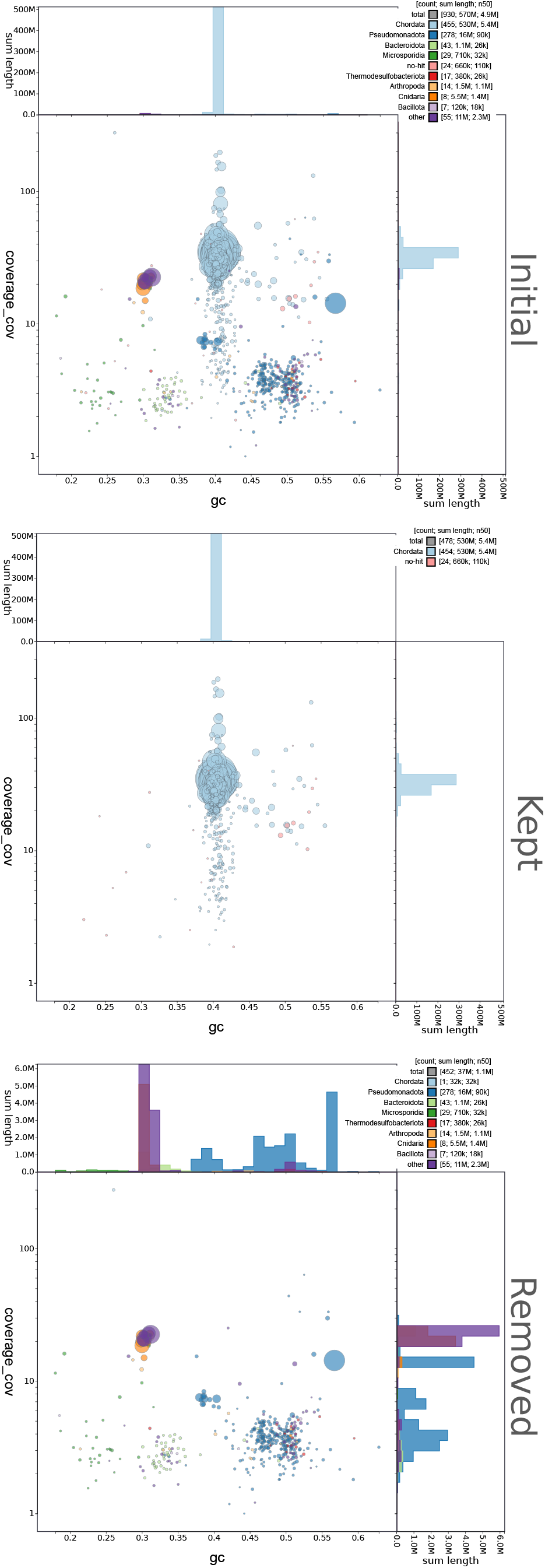
BlobPlots of the assembly of *B. schlosseri*. **Initial** corresponds to the results obtained with the primary assembly before scaffolding. **Kept** are the contigs kept for the scaffolding and **Removed** are those discarded as considered to be contamination.

**Fig. S5.**
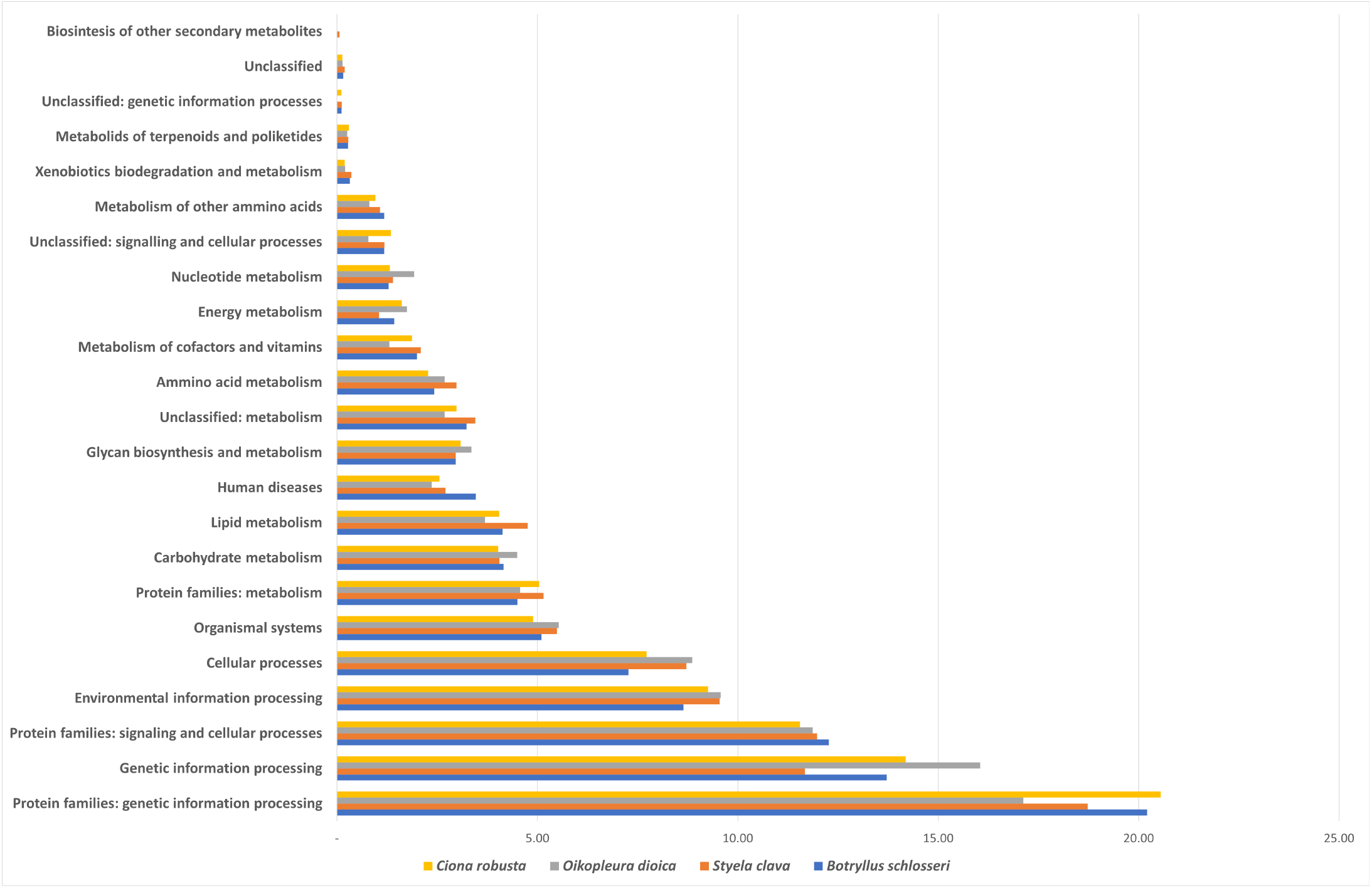
Comparison of the percentage of genes of *Botryllus schlosseri, Ciona robusta, Oikopleura dioica* and *Styela clava* assigned to different KEGG functional categories by BlastKOALA (44) .

**Fig. S6.**
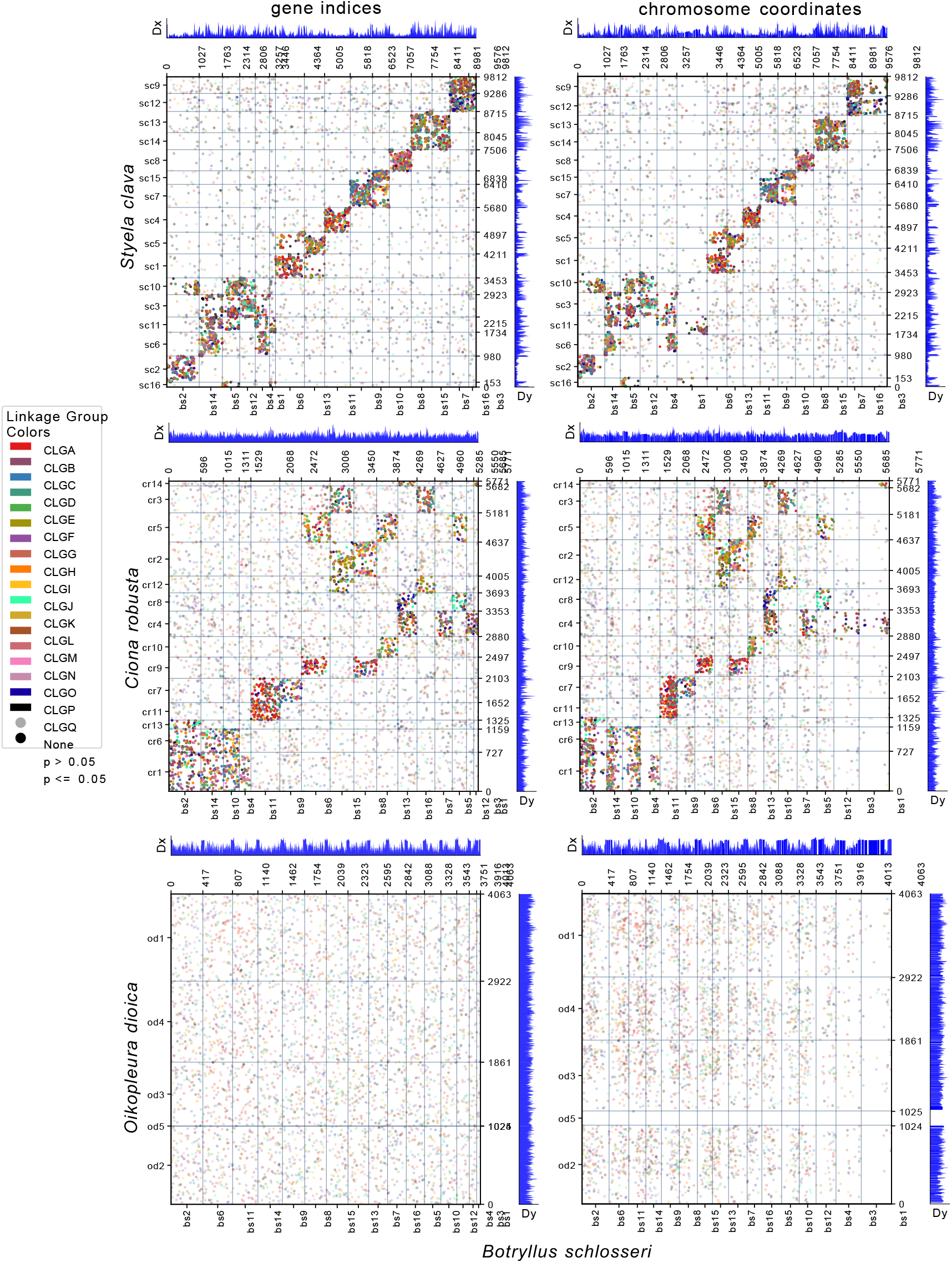
Investigation of synteny conservation among tunicate genomes. In the first column, dot plots depict the chromosome-scale scaffolds of *Botryllus schlosseri* (x-axis) plotted against those of *Styela clava, Ciona robusta* and *Oikopleura dioica* (y-axis). Each dot in the plot represents an ortholog, specifically a reciprocal best diamond blastp match between two species. The units of the x-and y-axes are the number of orthologous proteins: 9813, 5772 and 4064 orthologs found between the 16 *B. schlosseri* chromosome-scale scaffolds and the 16 of *S*.*plicata*, the 14 of *C*.*robusta* and the 5 of *O*.*dioica*, respectively. If there were chromosome breaks, Fisher’s exact test (FET) was used to calculate the significance of the interactions between the sub-chromosomal pieces. Otherwise, FET was calculated on whole chromosomes. The opacity of the dots depicts the significance of FET. Dots that are a solid color are in cells with a FET p-value less than or equal to 0.05. Dots that are translucent are in cells with a FET p-value greater than 0.05. Dx and Dy values allow to pinpoint places where there may be sudden breaks in synteny (49) The second column of the figure depicts the same information as the first one, but plotted following chromosome base pair coordinates rather than gene index. This is better suited for visualizing gene-poor regions of the chromosomes.

**Fig. S7.**
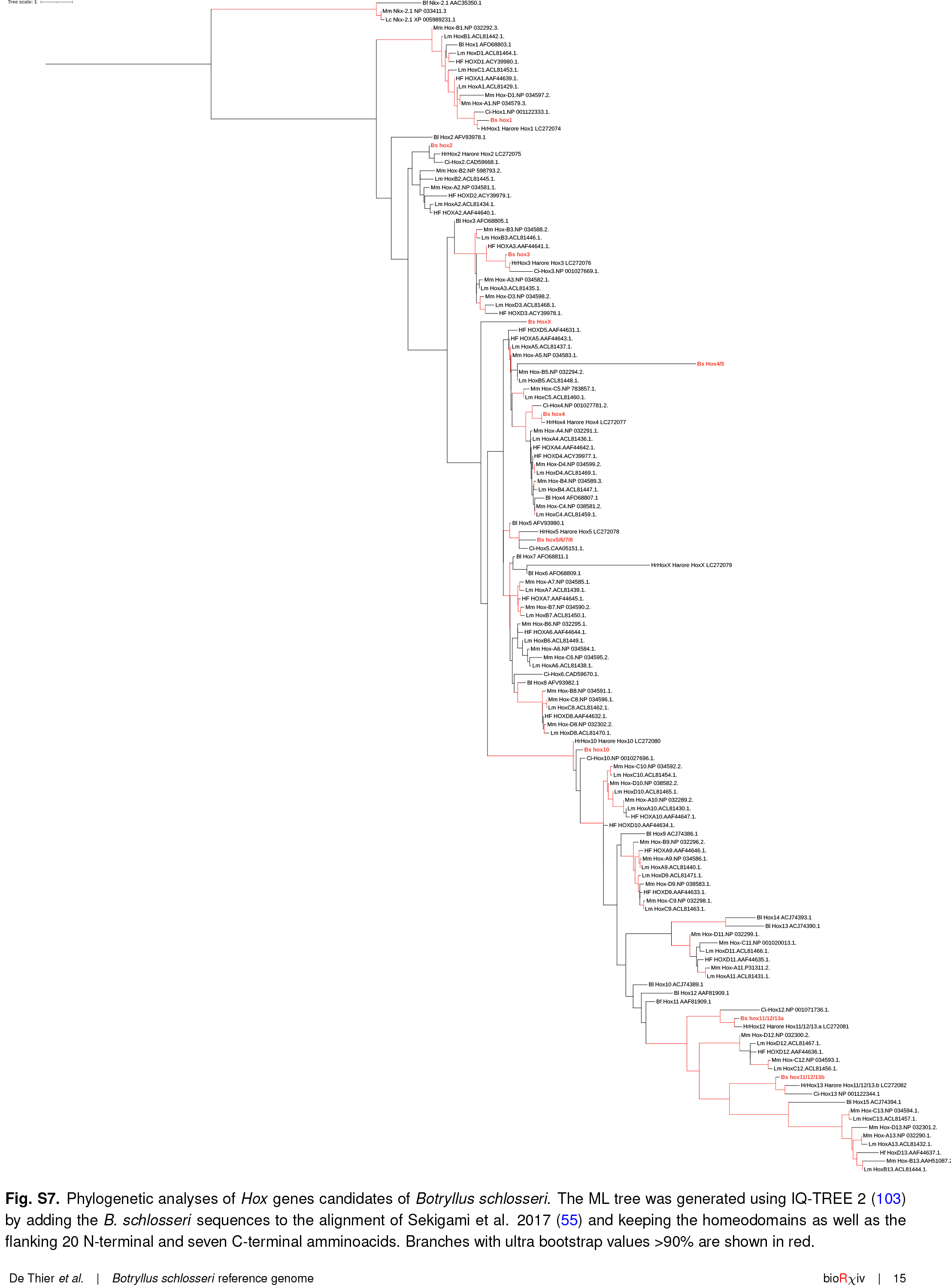
Phylogenetic analyses of *Hox* genes candidates of *Botryllus schlosseri*. The ML tree was generated using IQ-TREE 2 (103) by adding the *B. schlosseri* sequences to the alignment of Sekigami et al. 2017 (55) and keeping the homeodomains as well as the flanking 20 N-terminal and seven C-terminal amminoacids. Branches with ultra bootstrap values >90% are shown in red.

## Bibliography

1. Delsuc, F., Brinkmann, H., Chourrout, D. & Philippe, H. Tunicates and not cephalochordates are the closest living relatives of vertebrates. Nature 439, 965–968 (2006).

2. Alié, A., Hiebert, L. S., Scelzo, M. & Tiozzo, S. The eventful history of nonembryonic development in tunicates. Journal of Experimental Zoology Part B: Molecular and Developmental Evolution (2020).

3. Stolfi, A. & Brown, F. D. Tunicata. In Wanninger, A. (ed.) Evolutionary Developmental Biology of Invertebrates 6: Deuterostomia, 135–204 (Springer, Vienna, 2015).

4. Manni, L. et al. Ontology for the asexual development and anatomy of the colonial chordate Botryllus schlosseri. PLoS ONE 9, e96434 (2014).

5. Sabbadin, A., Zaniolo, G. & Majone, F. Determination of polarity and bilateral asymmetry in palleal and vascular buds of the ascidian Botryllus schlosseri. Developmental Biology 46, 79–87 (1975).

6. Nourizadeh, S., Kassmer, S., Rodriguez, D., Hiebert, L. S. & De Tomaso, A. W. Whole body regeneration and developmental competition in two botryllid ascidians. EvoDevo 12, 15 (2021-12-15).

7. Ricci, L. et al. The onset of whole-body regeneration in Botryllus schlosseri: morphological and molecular characterization. Frontiers in Cell and Developmental Biology 0, 173 (2022-02-14).

8. Laird, D. J., De Tomaso, A. W. & Weissman, I. L. Stem cells are units of natural selection in a colonial ascidian. Cell 123, 1351–1360 (2005).

9. Brown, F. D. et al. Early lineage specification of long-lived germline precursors in the colonial ascidian Botryllus schlosseri. Development 136, 3485–3494 (2009).

10. Laird, D. J. & De Tomaso, A. W. Predatory stem cells in the non-zebrafish chordate, Botryllus schlosseri. Zebrafish 1, 357–361 (2005).

11. Pancer, Z., Gershon, H. & Rinkevich, B. Coexistence and possible parasitism of somatic and germ cell lines in chimeras of the colonial urochordate Botryllus schlosseri. The Biological Bulletin 189, 106–112 (1995).

12. Stoner, D. S. & Weissman, I. L. Somatic and germ cell parasitism in a colonial ascidian: possible role for a highly polymorphic allorecognition system. Proceedings of the National Academy of Sciences of the United States of America 93, 15254–15259 (1996).

13. Manni, L. et al. Sixty years of experimental studies on the blastogenesis of the colonial tunicate Botryllus schlosseri. Developmental Biology 448, 293–308 (2019).

14. Kassmer, S. H., Rodriguez, D. & De Tomaso, A. W. Colonial ascidians as model organisms for the study of germ cells, fertility, whole body regeneration, vascular biology and aging. Current Opinion in Genetics & Development 39, 101–106 (2016).

15. Taketa, D. A. & De Tomaso, A. W. Botryllus schlosseri allorecognition: tackling the enigma. Developmental & Comparative Immunology 48, 254–265 (2015).

16. Nydam, M. L. Evolution of allorecognition in the tunicata. Biology 9, 1–13 (2020-06-01).

17. Epelbaum, A., Therriault, T. W., Paulson, A. & Pearce, C. M. Botryllid tunicates: Culture techniques and experimental procedures. Aquatic Invasions 4, 111–120 (2009).

18. Gasparini, F. et al. Sexual and asexual reproduction in the colonial ascidian Botryllus schlosseri. Genesis 53, 105–120 (2015).

19. Wawrzyniak, M. K., Matas Serrato, L. A. & Blanchoud, S. Long-term monitoring data logs of a recirculating artificial seawater based colonial ascidian aquaculture. Data in Brief 38, 107372 (2021-10).

20. Langenbacher, A. D., Rodriguez, D., Di Maio, A. & De Tomaso, A. W. Whole-mount fluorescent in situ hybridization staining of the colonial tunicate B otryllus schlosseri. genesis 53, 194–201 (2015).

21. Manni, L., Zaniolo, G., Cima, F., Burighel, P. & Ballarin, L. Botryllus schlosseri: A model ascidian for the study of asexual reproduction. Developmental Dynamics 236, 335–352 (2007).

22. Rodriguez, D. et al. Analysis of the basal chordate Botryllus schlosseri reveals a set of genes associated with fertility. BMC genomics 15, 1183 (2014).

23. Campagna, D. et al. Transcriptome dynamics in the asexual cycle of the chordate Botryllus schlosseri. BMC Genomics 17, 275 (2016).

24. Ricci, L. et al. Identification of differentially expressed genes from multipotent epithelia at the onset of an asexual development. Scientific Reports 6, 27357 (2016).

25. Rosental, B. et al. Complex mammalian-like haematopoietic system found in a colonial chordate. Nature 564, 425–429 (2018).

26. Kowarsky, M. et al. Sexual and asexual development: two distinct programs producing the same tunicate. Cell Reports 34, 108681 (2021).

27. Voskoboynik, A. et al. The genome sequence of the colonial chordate, Botryllus schlosseri. eLife 2, e00569 (2013).

28. Lawniczak, M. K. N. et al. Standards recommendations for the Earth BioGenome Project. Proceedings of the National Academy of Sciences 119, e2115639118 (2022).

29. De Tomaso, A. W., Saito, Y., Ishizuka, K. J., Palmeri, K. J. & Weissman, I. L. Mapping the genome of a model protochordate. I. A low resolution genetic map encompassing the fusion/histocompatibility (Fu/HC) locus of Botryllus schlosseri . Genetics 149, 277–287 (1998).

30. Cheng, H., Concepcion, G. T., Feng, X., Zhang, H. & Li, H. Haplotype-resolved de novo assembly using phased assembly graphs with hifiasm. Nature Methods 18, 170–175 (2021).

31. Li, K., Xu, P., Wang, J., Yi, X. & Jiao, Y. Identification of errors in draft genome assemblies at single-nucleotide resolution for quality assessment and improvement. Nature Communications 14, 6556 (2023).

32. Zhou, C., McCarthy, S. A. & Durbin, R. YaHS: yet another Hi-C scaffolding tool. Bioinformatics 39, btac808 (2023).

33. Bandi, V. et al. Visualization tools for genomic conservation. In Edwards, D. (ed.) Plant Bioinformatics: Methods and Protocols, 285–308 (Springer US, New York, NY, 2022).

34. Wang, Y. et al. MCScanX: a toolkit for detection and evolutionary analysis of gene synteny and collinearity. Nucleic Acids Research 40, e49 (2012).

35. Simão, F. A., Waterhouse, R. M., Ioannidis, P., Kriventseva, E. V. & Zdobnov, E. M. BUSCO: assessing genome assembly and annotation completeness with single-copy orthologs. Bioinformatics 31, 3210–3212 (2015).

36. Guiglielmoni, N., Houtain, A., Derzelle, A., Van Doninck, K. & Flot, J.-F. Overcoming uncollapsed haplotypes in long-read assemblies of non-model organisms. BMC Bioinformatics 22, 303 (2021).

37. Simion, P. et al. Chromosome-level genome assembly reveals homologous chromosomes and recombination in asexual rotifer Adineta vaga. Science Advances 7, eabg4216 (2021).

38. Gabriel, L. et al. BRAKER3: Fully automated genome annotation using RNA-Seq and protein evidence with GeneMark-ETP, AUGUSTUS and TSEBRA. bioRxiv 2023.06.10.544449 (2023).

39. Haas, B. J. Improving the Arabidopsis genome annotation using maximal transcript alignment assemblies. Nucleic Acids Research 31, 5654–5666 (2003).

40. Haas, B. J. et al. Automated eukaryotic gene structure annotation using EVidenceModeler and the Program to Assemble Spliced Alignments. Genome Biology 9, R7 (2008).

41. Cantalapiedra, C. P., Hernández-Plaza, A., Letunic, I., Bork, P. & Huerta-Cepas, J. eggNOG-mapper v2: functional annotation, orthology assignments, and domain prediction at the metagenomic scale. Molecular Biology and Evolution 38, 5825–5829 (2021).

42. Blum, M. et al. The InterPro protein families and domains database: 20 years on. Nucleic Acids Research 49, D344–D354 (2021).

43. Jones, P. et al. InterProScan 5: genome-scale protein function classification. Bioinformatics 30, 1236–1240 (2014).

44. Kanehisa, M., Sato, Y. & Morishima, K. BlastKOALA and GhostKOALA: KEGG tools for functional characterization of genome and metagenome sequences. Journal of Molecular Biology 428, 726–731 (2016).

45. Wei, J. et al. Genomic basis of environmental adaptation in the leathery sea squirt (Styela clava). Molecular Ecology Resources 20, 1414–1431 (2020).

46. Satou, Y. et al. A nearly complete genome of Ciona intestinalis type A (C. robusta) reveals the contribution of inversion to chromosomal evolution in the genus Ciona. Genome Biology and Evolution 11, 3144–3157 (2019).

47. Bliznina, A. et al. Telomere-to-telomere assembly of the genome of an individual Oikopleura dioica from Okinawa using Nanopore-based sequencing. BMC Genomics 22, 222 (2021).

48. Delsuc, F. et al. A phylogenomic framework and timescale for comparative studies of tunicates. BMC Biology 16, 1–14 (2018).

49. Simakov, O. et al. Deeply conserved synteny resolves early events in vertebrate evolution. Nature Ecology & Evolution 4, 820–830 (2020-06).

50. Schultz, D. T. et al. Ancient gene linkages support ctenophores as sister to other animals. Nature 618, 110–117 (2023).

51. Plessy, C. et al. Extreme genome scrambling in marine planktonic Oikopleura dioica cryptic species. Genome Research (2024).

52. Simakov, O. et al. Deeply conserved synteny and the evolution of metazoan chromosomes. Science Advances 8, eabi5884 (2022).

53. Monteiro, A. S. & Ferrier, D. E. Hox genes are not always Colinear. International Journal of Biological Sciences 95–103 (2006).

54. DeBiasse, M. B., Colgan, W. N., Harris, L., Davidson, B. & Ryan, J. F. Inferring tunicate relationships and the evolution of the tunicate Hox cluster with the genome of Corella inflata. Genome Biology and Evolution 12, 948–964 (2020).

55. Sekigami, Y. et al. Hox gene cluster of the ascidian, Halocynthia roretzi, reveals multiple ancient steps of cluster disintegration during ascidian evolution. Zoological Letters 3, 17 (2017).

56. Gaunt, S. J. Seeking sense in the Hox gene cluster. Journal of Developmental Biology 10, 48 (2022).

57. Blanchoud, S., Rutherford, K., Zondag, L., Gemmell, N. J. & Wilson, M. J. De novo draft assembly of the Botrylloides leachii genome provides further insight into tunicate evolution. Scientific Reports 8, 5518 (2018).

58. Sim, S. B., Corpuz, R. L., Simmonds, T. J. & Geib, S. M. HiFiAdapterFilt, a memory efficient read processing pipeline, prevents occurrence of adapter sequence in PacBio HiFi reads and their negative impacts on genome assembly. BMC Genomics 23, 157 (2022).

59. Bolger, A. M., Lohse, M. & Usadel, B. Trimmomatic: a flexible trimmer for Illumina sequence data. Bioinformatics 30, 2114–2120 (2014).

60. Andrews, S. FastQC: A quality control tool for high throughput sequence data. Available online at: http://www.bioinformatics.babraham.ac.uk/projects/fastqc/ (2010).

61. Wang, L. et al. Genome assembly and annotation of Periplaneta americana reveal a comprehensive cockroach allergen profile. Allergy 78, 1088–1103 (2023).

62. Nowoshilow, S. et al. The axolotl genome and the evolution of key tissue formation regulators. Nature 554, 50–55 (2018).

63. Schneider, C. A., Rasband, W. S. & Eliceiri, K. W. NIH Image to ImageJ: 25 years of image analysis. Nature Methods 9, 671–675 (2012).

64. Kokot, M., Dlugosz, M. & Deorowicz, S. KMC 3: counting and manipulating k-mer statistics. Bioinformatics 33, 2759–2761 (2017).

65. Ranallo-Benavidez, T. R., Jaron, K. S. & Schatz, M. C. GenomeScope 2.0 and Smudgeplot for reference-free profiling of polyploid genomes. Nature Communications 11, 1432 (2020).

66. Huang, S., Kang, M. & Xu, A. HaploMerger2: rebuilding both haploid sub-assemblies from high-heterozygosity diploid genome assembly. Bioinformatics 33, 2577–2579 (2017).

67. Challis, R., Richards, E., Rajan, J., Cochrane, G. & Blaxter, M. BlobToolKit – interactive quality assessment of genome assemblies. G3: Genes|Genomes|Genetics 10, 1361–1374 (2020).

68. Camacho, C. et al. BLAST+: architecture and applications. BMC Bioinformatics 10, 421 (2009).

69. Buchfink, B., Reuter, K. & Drost, H.-G. Sensitive protein alignments at tree-of-life scale using DIAMOND. Nature Methods 18, 366–368 (2021).

70. Li, H. New strategies to improve minimap2 alignment accuracy. Bioinformatics 37, 4572–4574 (2021).

71. Ghurye, J. et al. Integrating Hi-C links with assembly graphs for chromosome-scale assembly. PLOS Computational Biology 15, e1007273 (2019).

72. Harry, E. PretextMap (Paired REad TEXTure Mapper): Converts sam formatted read pairs into genome contact maps. https://github.com/wtsi-hpag/pretextmap.

73. Harry, E. PretextView (Paired REad TEXTure Viewer): A desktop application for viewing pretext contact maps. https://github.com/wtsi-hpag/pretextview.

74. Shen, W., Le, S., Li, Y. & Hu, F. SeqKit: a cross-platform and ultrafast toolkit for FASTA/Q file manipulation. PLoS ONE 11, e0163962 (2016).

75. Mapleson, D., Garcia Accinelli, G., Kettleborough, G., Wright, J. & Clavijo, B. J. KAT: a K-mer analysis toolkit to quality control NGS datasets and genome assemblies. Bioinformatics 33, 574–576 (2017).

76. Manni, M., Berkeley, M. R., Seppey, M., Simão, F. A. & Zdobnov, E. M. BUSCO update: novel and streamlined workflows along with broader and deeper phylogenetic coverage for scoring of eukaryotic, prokaryotic, and viral genomes. Molecular Biology and Evolution 38, 4647–4654 (2021).

77. Flynn, J. M. et al. RepeatModeler2 for automated genomic discovery of transposable element families. Proceedings of the National Academy of Sciences 117, 9451–9457 (2020).

78. Smit, A., Hubley, R. & Green, P. RepeatMasker Open-4.0. 2013-2015 <http://www.repeatmasker.org>.

79. Dobin, A. et al. STAR: ultrafast universal RNA-seq aligner. Bioinformatics 29, 15–21 (2013).

80. Kuznetsov, D. et al. OrthoDB v11: annotation of orthologs in the widest sampling of organismal diversity. Nucleic Acids Research 51, D445–D451 (2023).

81. Lomsadze, A. Gene identification in novel eukaryotic genomes by self-training algorithm. Nucleic Acids Research 33, 6494–6506 (2005).

82. Stanke, M., Schöffmann, O., Morgenstern, B. & Waack, S. Gene prediction in eukaryotes with a generalized hidden Markov model that uses hints from external sources. BMC Bioinformatics 7, 62 (2006).

83. Lomsadze, A., Burns, P. D. & Borodovsky, M. Integration of mapped RNA-Seq reads into automatic training of eukaryotic gene finding algorithm. Nucleic Acids Research 42, e119–e119 (2014).

84. Gotoh, O. A space-efficient and accurate method for mapping and aligning cDNA sequences onto genomic sequence. Nucleic Acids Research 36, 2630–2638 (2008).

85. Iwata, H. & Gotoh, O. Benchmarking spliced alignment programs including Spaln2, an extended version of Spaln that incorporates additional species-specific features. Nucleic Acids Research 40, e161–e161 (2012).

86. Buchfink, B., Xie, C. & Huson, D. H. Fast and sensitive protein alignment using DIAMOND. Nature Methods 12, 59–60 (2015).

87. Brůna, T., Lomsadze, A. & Borodovsky, M. GeneMark-EP+: eukaryotic gene prediction with self-training in the space of genes and proteins. NAR Genomics and Bioinformatics 2, lqaa026 (2020).

88. Pertea, G. & Pertea, M. GFF Utilities: GffRead and GffCompare. F1000Research 9, ISCB Comm J–304 (2020).

89. Kovaka, S. et al. Transcriptome assembly from long-read RNA-seq alignments with StringTie2. Genome Biology 20, 278 (2019).

90. Stanke, M., Diekhans, M., Baertsch, R. & Haussler, D. Using native and syntenically mapped cDNA alignments to improve de novo gene finding. Bioinformatics 24, 637–644 (2008).

91. Hoff, K. J., Lomsadze, A., Borodovsky, M., Stanke, M. & Kollmar, M. Whole-genome annotation with BRAKER. In Gene prediction: methods and protocols, no. 1962 in Methods in Molecular Biology, 65–95 (Springer, 2019).

92. Hoff, K. J., Lange, S., Lomsadze, A., Borodovsky, M. & Stanke, M. BRAKER1: unsupervised RNA-Seq-based genome annotation with GeneMark-ET and AUGUSTUS. Bioinformatics 32, 767–769 (2016).

93. Brůna, T., Hoff, K. J., Lomsadze, A., Stanke, M. & Borodovsky, M. BRAKER2: automatic eukaryotic genome annotation with GeneMark-EP+ and AUGUSTUS supported by a protein database. NAR Genomics and Bioinformatics 3, lqaa108 (2021).

94. Pertea, M. et al. StringTie enables improved reconstruction of a transcriptome from RNA-seq reads. Nature Biotechnology 33, 290–295 (2015).

95. Haas, B. TransDecoder https://github.com/TransDecoder/TransDecoder.

96. Huerta-Cepas, J. et al. eggNOG 5.0: a hierarchical, functionally and phylogenetically annotated orthology resource based on 5090 organisms and 2502 viruses. Nucleic Acids Research 47, D309–D314 (2019).

97. Palmer, J. M. & Stajich, J. E. Funannotate (2023). URL https://github.com/nextgenusfs/funannotate. Original-date: 2015-12-18T20:21:15Z.

98. Dardaillon, J. et al. ANISEED 2019: 4D exploration of genetic data for an extended range of tunicates. Nucleic Acids Research 48, D668–D675 (2020).

99. Castellano, K. R., Batta-Lona, P., Bucklin, A. & O’Neill, R. J. Salpa genome and developmental transcriptome analyses reveal molecular flexibility enabling reproductive success in a rapidly changing environment. Scientific Reports 13, 21056 (2023).

100. Dainat, J. AGAT: Another Gff Analysis Toolkit to handle annotations in any GTF/GFF format. (Version v0.7.0). Zenodo. 10.5281/zenodo.3552717.

101. Edgar, R. C. MUSCLE: multiple sequence alignment with high accuracy and high throughput. Nucleic Acids Research 32, 1792–1797 (2004).

102. Larsson, A. AliView: a fast and lightweight alignment viewer and editor for large datasets. Bioinformatics 30, 3276–3278 (2014).

103. Minh, B. Q. et al. IQ-TREE 2: new models and efficient methods for phylogenetic inference in the genomic era. Molecular Biology and Evolution 37, 1530–1534 (2020).

104. Jones, D. T., Taylor, W. R. & Thornton, J. M. The rapid generation of mutation data matrices from protein sequences. Bioinformatics 8, 275–282 (1992).

105. Yang, Z. A space-time process model for the evolution of DNA sequences. Genetics 139, 993–1005 (1995).

106. Kalyaanamoorthy, S., Minh, B. Q., Wong, T. K. F., von Haeseler, A. & Jermiin, L. S. ModelFinder: fast model selection for accurate phylogenetic estimates. Nature Methods 14, 587–589 (2017).

107. Schwarz, G. Estimating the dimension of a model. The Annals of Statistics 6, 461–464 (1978).

108. Hoang, D. T., Chernomor, O., von Haeseler, A., Minh, B. Q. & Vinh, L. S. UFBoot2: improving the ultrafast bootstrap approximation. Molecular Biology and Evolution 35, 518–522 (2018).

